# Glycerol improves the viability of a cryopreserved choanoflagellate

**DOI:** 10.1101/2024.10.02.616207

**Authors:** Sheel Chandra, Florentine U. Rutaganira

**Affiliations:** Howard Hughes Medical Institute and the Department of Molecular and Cell Biology, University of California, Berkeley, CA, 94720; Department of Biology, University of Pennsylvania, Philadelphia, PA, 19104; Departments of Biochemistry and Developmental Biology, Stanford University School of Medicine, Stanford, CA 94305

**Keywords:** choanoflagellate, dimethyl sulfoxide, glycerol, cryopreservation, protocol, protist, bacteria, co-culture, liquid phase nitrogen, ultra-low temperature freezer

## Abstract

The colonial choanoflagellate *Salpingoeca rosetta* is a tractable model system for studying the origins of multicellularity, but long-term storage strategies for this species have not been tested. In this study, we probed each stage of cryopreservation (freeze-down, long-term storage, recovery) to identify the optimal protocol for recovery of *S. rosetta* and co-cultured bacterial cells. Dimethyl sulfoxide (Me2SO; commonly referred to as DMSO), the current cryoprotective agent (CPA) standard, proved to be worse than glycerol at comparable concentrations. Samples treated with either CPA at 5% showed the poorest recovery. Our results identified 15% glycerol as the most effective CPA for both *S. rosetta* and *Echinicola pacifica*. We also determined that ultra-low temperature freezers can be sufficient for short-term storage. We propose 15% glycerol and liquid phase nitrogen as the standard cryopreservation protocol for *S. rosetta* cultures and as a starting point for testing long-term storage strategies for other choanoflagellates and heterotrophic protists.

## 1. Brief Communication

Choanoflagellates are heterotrophic, single-celled eukaryotes and the closest living relatives of animals [1–3]. The colony-forming choanoflagellate species *Salpingoeca rosetta*, an emerging model system, presents a valuable resource for elucidating the basis of multicellular development in animals [4]. To reduce the need for continuous field sampling, cultures require long-term storage for standardizing and generating sustainable cell stocks. Because choanoflagellates are co-cultured with bacteria (their prey), cryopreservation conditions must be suitable for both organisms [5]. The only reported cryopreservation protocol for choanoflagellates was validated with *Monosiga brevicollis* [5], and long-term storage strategies specific to *S. rosetta* remain unexplored.

Cryopreservation protocols typically follow three stages [6]. The first stage is freeze-down, which involves the addition of cryoprotectant agents (CPAs) and rate-controlled cooling. Freeze-down presents potential threats to cellular integrity [7]. Rapid cooling can cause intracellular ice formation that is usually fatal. Slow cooling triggers extracellular ice formation, with cellular dehydration, exposure to high solute concentrations, and eventual osmotic shock. To mitigate cryoinjury and promote optimal cooling, CPAs such as dimethyl sulfoxide (Me2SO) and glycerol are commonly used, but high concentrations can be cytotoxic. Thus, an appropriate CPA is an important factor in protocols intended to optimize survival (Figure 1A). The second stage of cryopreservation is long-term storage, with transfer of frozen samples to indefinite storage at -196°C (either in a liquid nitrogen dewar or cryogenic freezer), and the third stage is recovery, which includes the thawing and culturing of samples.

**Figure 1.**
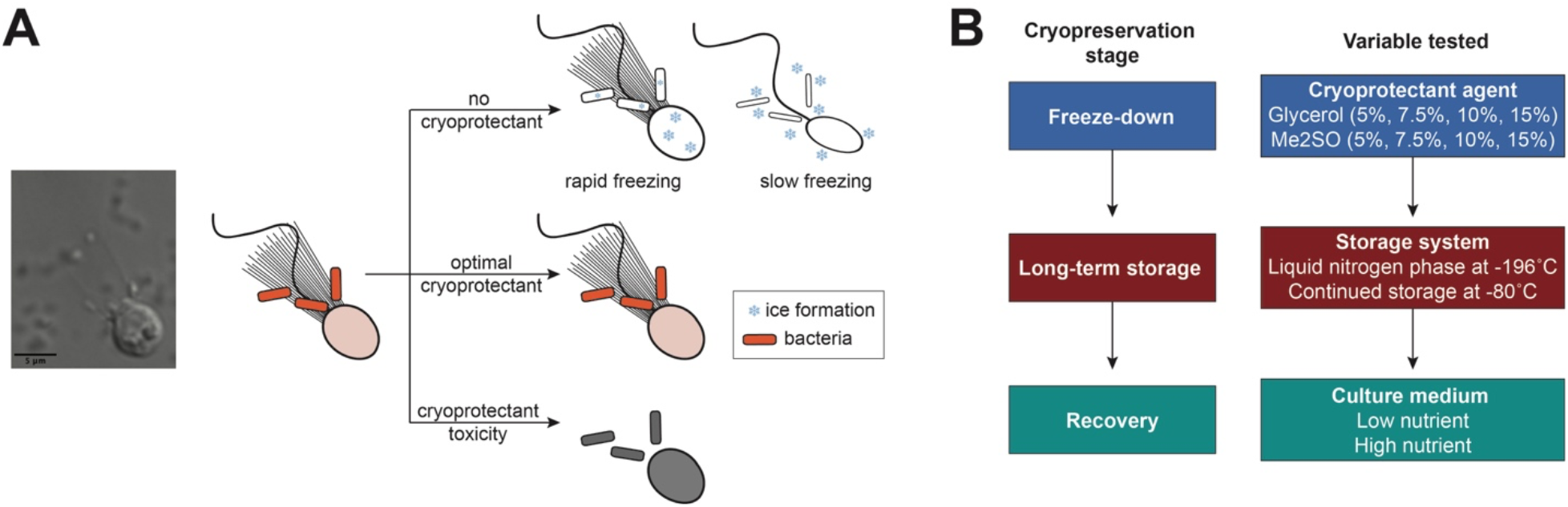
Rationales and procedures for *S. rosetta* cryopreservation strategies. (A) Choanoflagellates are bacteriovores and are co-cultured with their prey. The freeze-down of samples without a cryoprotectant agent (CPA) can result in cryoinjury: rapid freezing can lead to fatal intracellular ice formation while slow cooling can trigger extracellular ice formation, which eventually results in osmotic shock. Excess concentrations of CPAs such as Me2SO and glycerol can be toxic to the cell. The optimal cryoprotectant mitigates freeze-down injuries while causing the least amount of toxicity to the cell. (B) Schematic describing the three stages of a standard cryopreservation protocol and the variables at each stage that we tested in this study.

Here, to determine the optimal protocol for preserving *S. rosetta* cultures (Figure 1A), we tested the relevant variables for each stage in three sets of experiments (Figure 1B, Table S1). We chose 10% Me2SO (Millipore Sigma, Cat# D2650; or GoldBio, Cat# D-361-10) as a starting point because of its efficiency in preserving *M. brevicollis* stocks [5] and its use as a standard CPA for mammalian cell lines [8]. Because *S. rosetta* is co-cultured with its prey bacterium, *Echinicola pacifica*, we also assessed glycerol (Fisher Scientific, Cat# G33500), as it is the standard CPA for bacterial cultures [9].

Each CPA was added to cultures at maximal density (∼10^6^ cells/mL) in 1 mL cryogenic tubes with internal threading (Nunc™, Thermo Scientific, Cat# 366656), prepared in a biosafety cabinet. Within 10 minutes, cryovials were placed in a Mr. Frosty™ (Thermo Scientific, Cat# 5100-0001) or FreezeCell™ (Thomas Scientific, Cat# 1156P21) controlled-rate freezer (−1°C/min) and settled into a -80°C ultra-low temperature freezer for at least 24 hours. For cultures cryopreserved at -196°C, tubes were transferred to cryogenic boxes that were placed in a cryogenic dewar with the boxes submerged in liquid nitrogen (LN2) and cryopreserved for different storage lengths (Table S1). To recover cultures, tubes were thawed in a 37°C water bath for ∼1 minute (when tube contents had thawed), and the thawed cells were transferred to a T25 flask containing 9 or 14 mL of culture media in a biosafety cabinet.

We first compared the effects of 10% Me2SO versus 10% glycerol on cell viability in *S. rosetta* recovery, using samples from both conditions that were maintained at -196°C in a LN2 cryogenic dewar for less than a month (Table S1). At 96 hours after thawing, samples in 10% glycerol had the higher recovered mean cell density (Figure 2A). As a more direct test of cell viability, we also compared cell recovery between 10% Me2SO and 10% glycerol control samples that were frozen-down but not maintained in long-term storage (Figure 2B). Both control groups showed lower mean cell density compared to samples recovered from long-term storage, suggesting that direct thawing of cells after overnight storage at -80°C could slow down cell recovery. The 10% glycerol control still had the higher mean cell density compared with 10% Me2SO (Figure 2B), suggesting reduced toxic effects on cell viability with glycerol.

**Figure 2.**
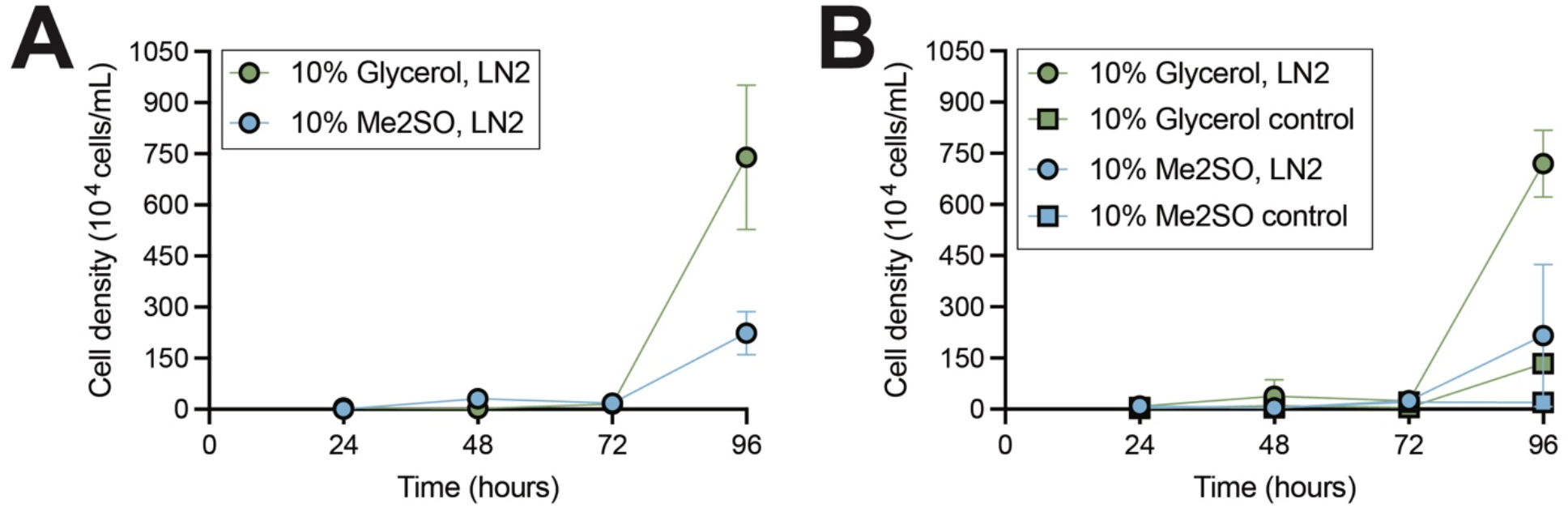
Glycerol outperforms Me2SO in preserving *S. rosetta* cultures. (A-B) *S. rosetta* recovery, measured as cell density (10,000 cells/mL) in samples cultured at 24, 48, 72, and 96 hours post-thawing after storage in a liquid nitrogen cryogenic (LN2) dewar for (A) biological replicate 1, which has two technical replicates in 10% glycerol and 10% Me2SO, and (B) biological replicate 2, which has three technical replicates in 10% glycerol and 10% Me2SO, and one replicate each in 10% Me2SO and 10% glycerol as control. Mean density in each condition is plotted ± standard deviation.

Because 10% Me2SO underperformed against 10% glycerol in cell recovery, we tested whether other Me2SO concentrations might yield better recovery. Initial tests showed no significant difference in viability among samples preserved in 5%, 7.5%, or 10% Me2SO (Figure S1A, Table S1), but a second experiment comparing 5%, 10%, and 15% Me2SO showed poor recovery with 5% Me2SO, even at 145 hours post-thawing (Figure S2A, Table S1). The 15% Me2SO samples showed no significantly improved cell proliferation compared to 10% Me2SO (Figure S2A).

Our finding of negative effects of overnight (short-term) storage at -80°C led us to test recovery of cells cryopreserved with Me2SO and stored at -80°C long term. Compared with storage in LN2, cells cryopreserved with 15% Me2SO were less viable when stored at -80°C (Figures S1B, S2B). The results indicated that 10% Me2SO and storage in LN2, the current CPA and storage standard, is optimal when using Me2SO as a CPA. However, 10% Me2SO underperformed 10% glycerol in cell recovery (Figure 2A). These results motivated us to investigate glycerol as a potential CPA standard in the long-term preservation of *S. rosetta* cultures.

We found that *S. rosetta* cultures preserved in 10% glycerol had higher cell densities when recovered in high-compared with low-nutrient media (Figure 3A), so all subsequent experiments were conducted using high-nutrient media (Table S1). We preserved cultures with 5%, 10%, or 15% glycerol to identify the optimal concentration and found that preservation with 15% glycerol yielded faster cell recovery by 100 hours post-thaw in LN2 (Figure 3B). Decreasing the glycerol concentration did not improve cell recovery (Figures 3B, S1C).

**Figure 3.**
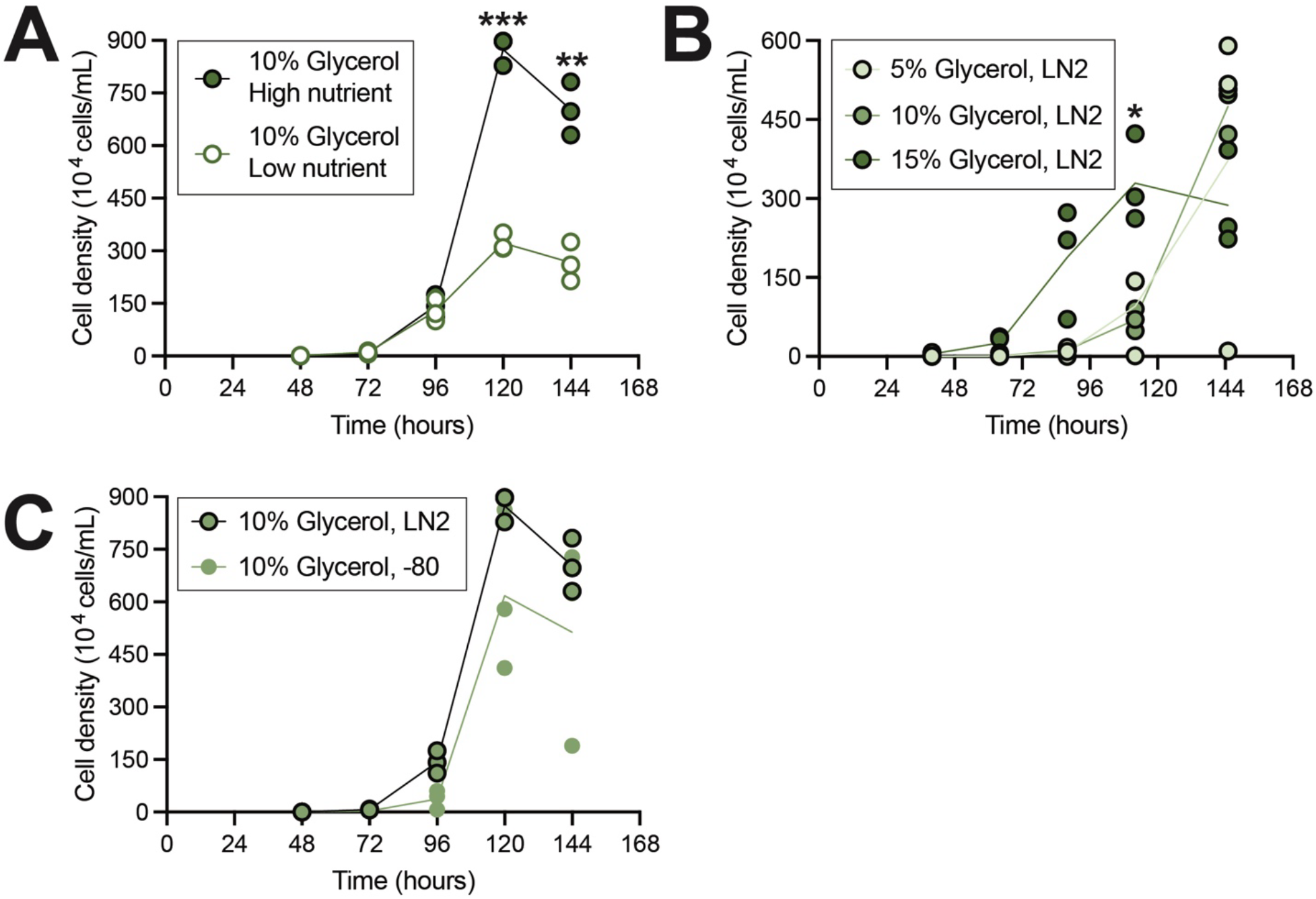
*S. rosetta* cultures were most viable when preserved with 15% glycerol and recovered in high-nutrient media. (A) *S. rosetta* cultures cryopreserved with 10% glycerol and recovered in high-nutrient media were more viable than cultures recovered in low-nutrient media. At 120 and 144 hours, cell densities of cultures recovered in high-nutrient media were significantly higher compared with recovery from cultures from low-nutrient media (**p=0.0094; ***p=0.007). (B) *S. rosetta* cultures cryopreserved in 15% glycerol, stored in LN2, and recovered in high-nutrient media were more viable compared to cryopreservation in 5% or 10% glycerol. At 112 hours, *S. rosetta* cultures preserved in 15% glycerol showed the best viability post-storage in LN2 (*p = 0.0545 for 5% vs 15% glycerol; p=0.0505 for 10% vs 15% glycerol. (C) *S. rosetta* cultures cryopreserved with 10% glycerol and recovered in high-nutrient media were equally viable with storage in LN2 or at -80°C. Significance was determined by a two-way ANOVA multiple comparisons test with Geisser–Greenhouse and Šídák corrections (calculations and charts, Prism v. 10.1.1).

To examine the effects of different long-term storage methods on the efficacy of glycerol, we stored glycerol-preserved samples in LN2 or in an ultra-low temperature freezer at -80°C. With either storage method, we observed the slowest recovery with 5% glycerol (Figures 3B, S1C, S1E, S1F). Samples preserved in 10% glycerol showed better recovery when stored in LN2 compared with -80°C but the results were not significant (Figure 3C). The cell density in 15% glycerol–preserved samples was comparable between those stored at LN2 and at -80°C (Figure S1D), indicating negligible differences in cell viability between these methods post-thawing if glycerol is used as a CPA at 10% or 15%. Our results for *S. rosetta* overall show that 15% glycerol is superior to other glycerol concentrations and to the current CPA standard, 10% Me2SO. In general, glycerol-treated samples showed higher growth, faster recovery, and less variability than Me2SO-preserved samples (Figures S3A, S3B).

Previous studies have documented growth inhibition in certain bacterial cultures treated with Me2SO [9]. In our experiments with *E. pacifica*, which are usually co-cultured with *S. rosetta*, we found that glycerol was better than Me2SO for maintaining the colonies (Figure S3C). Although Me2SO enters cells more rapidly than glycerol, it is much more toxic [7,10]. *S. rosetta* and *E. pacifica* cells likely exhibited reduced recovery with Me2SO because of this toxicity.

Assessment of other factors to optimize choanoflagellate cryopreservation was beyond the scope of our study. Although lower Me2SO concentrations did not improve cell viability, maintaining cell suspensions at low temperatures before freezing might reduce exposure to Me2SO and mitigate its toxicity [7]. Me2SO and glycerol are the most commonly used CPAs, but other permeating CPAs (e.g., ethylene glycol, propylene glycol) and non-permeating CPAs (e.g., trehalose) are less toxic than Me2SO and might better support choanoflagellate and bacterial recovery [7,9]. With our optimal CPA, 15% glycerol, we found similar recoveries between storage at -80°C and storage at LN2, but experiments with longer storage periods (>1 year) are needed to ascertain whether choanoflagellate cultures can always be recovered from - 80°C storage. Pending such studies, we conclude that LN2 storage is optimal for periods longer than a year.

Finally, here we focused our efforts on identifying optimal cryopreservation conditions for *S. rosetta*, a marine craspedid choanoflagellate. Choanoflagellates with different biology (e.g., freshwater, loricate) may have unique cryopreservation requirements, but for determining optimal conditions, we recommend 15% glycerol and storage in LN2 as an initial approach.

## 2. Funding information

This work was supported by the Howard Hughes Medical Institute (HHMI) Investigator and Hanna Gray programs.

### CRediT authorship contribution statement

**Sheel Chandra**: Conceptualization, Methodology, Formal Analysis, Investigation, Writing - original draft, Writing - Review & Editing. **Florentine U. Rutaganira**: Conceptualization, Methodology, Validation, Formal Analysis, Investigation, Resources, Writing - original draft, Writing - Review & Editing, Visualization, Supervision, Project Administration, Funding acquisition

## Acknowledgments

We are grateful for the support of Professor Nicole King (HHMI and the Department of Molecular and Cell Biology, University of California, Berkeley), her research group, and the broader choanoflagellate community.

## References

[1] N. King, M.J. Westbrook, S.L. Young, A. Kuo, M. Abedin, J. Chapman, S. Fairclough, U. Hellsten, Y. Isogai, I. Letunic, M. Marr, D. Pincus, N. Putnam, A. Rokas, K.J. Wright, R. Zuzow, W. Dirks, M. Good, D. Goodstein, D. Lemons, W. Li, J.B. Lyons, A. Morris, S. Nichols, D.J. Richter, A. Salamov, J.G.I. Sequencing, P. Bork, W.A. Lim, G. Manning, W.T. Miller, W. McGinnis, H. Shapiro, R. Tjian, I.V. Grigoriev, D. Rokhsar, The genome of the choanoflagellate Monosiga brevicollis and the origin of metazoans, Nature 451 (2008) 783– 788.

[2] D.S. Booth, H. Szmidt-Middleton, N. King, Transfection of choanoflagellates illuminates their cell biology and the ancestry of animal septins, Mol. Biol. Cell 29 (2018) 3026–3038.

[3] D.S. Booth, N. King, Genome editing enables reverse genetics of multicellular development in the choanoflagellate Salpingoeca rosetta, Elife 9 (2020). 10.7554/eLife.56193.

[4] M.J. Dayel, R.A. Alegado, S.R. Fairclough, T.C. Levin, S.A. Nichols, K. McDonald, N. King, Cell differentiation and morphogenesis in the colony-forming choanoflagellate Salpingoeca rosetta, Dev. Biol. 357 (2011) 73–82.

[5] N. King, S.L. Young, M. Abedin, M. Carr, B.S.C. Leadbeater, Long-term frozen storage of choanoflagellate cultures, Cold Spring Harb. Protoc. 2009 (2009) db.prot5149.

[6] T.H. Jang, S.C. Park, J.H. Yang, J.Y. Kim, J.H. Seok, U.S. Park, C.W. Choi, S.R. Lee, J. Han, Cryopreservation and its clinical applications, Integr. Med. Res. 6 (2017) 12–18.

[7] K.A. Murray, M.I. Gibson, Chemical approaches to cryopreservation, Nat. Rev. Chem. 6 (2022) 579–593.

[8] D.E. Pegg, The history and principles of cryopreservation, Semin. Reprod. Med. 20 (2002) 5–13.

[9] Z. Hubálek, Protectants used in the cryopreservation of microorganisms, Cryobiology 46 (2003) 205–229.

[10] J. Galvao, B. Davis, M. Tilley, E. Normando, M.R. Duchen, M.F. Cordeiro, Unexpected low-dose toxicity of the universal solvent DMSO, FASEB J. 28 (2014) 1317–1330.

