## Supplementary Information for "Glycerol improves the viability of a cryopreserved choanoflagellate"

**Supplemental Table 1.** Experimental conditions for cryopreservation testing

| | Experiment 1 (Figures 2, S3C) | | Experiment 2 (Figures S1, S3) | | Experiment 3 (Figures 3, S1–S3) | |
|---|---|---|---|---|---|---|
| **# of cells cryopreserved** | ~4 million | | 10 million | | 2.5 million | |
| **Nutrient media** | High nutrient | | High nutrient | Low nutrient | High nutrient | |
| **Cryoprotectant** | 10% Me2SO | 10% glycerol | 5% Me2SO<br>7.5% Me2SO<br>10% Me2SO | 5% glycerol<br>7.5% glycerol<br>10% glycerol | 5% Me2SO<br>10% Me2SO<br>15% Me2SO | 5% glycerol<br>10% glycerol<br>15% glycerol |
| **Cold storage** | Liquid nitrogen | | Liquid nitrogen | -80°C freezer | Liquid nitrogen | -80°C freezer |
| **Storage period** | Biological rep. 1: 2 days<br>Biological rep. 2: 2 weeks | | 14 months | | 6.5 months | |

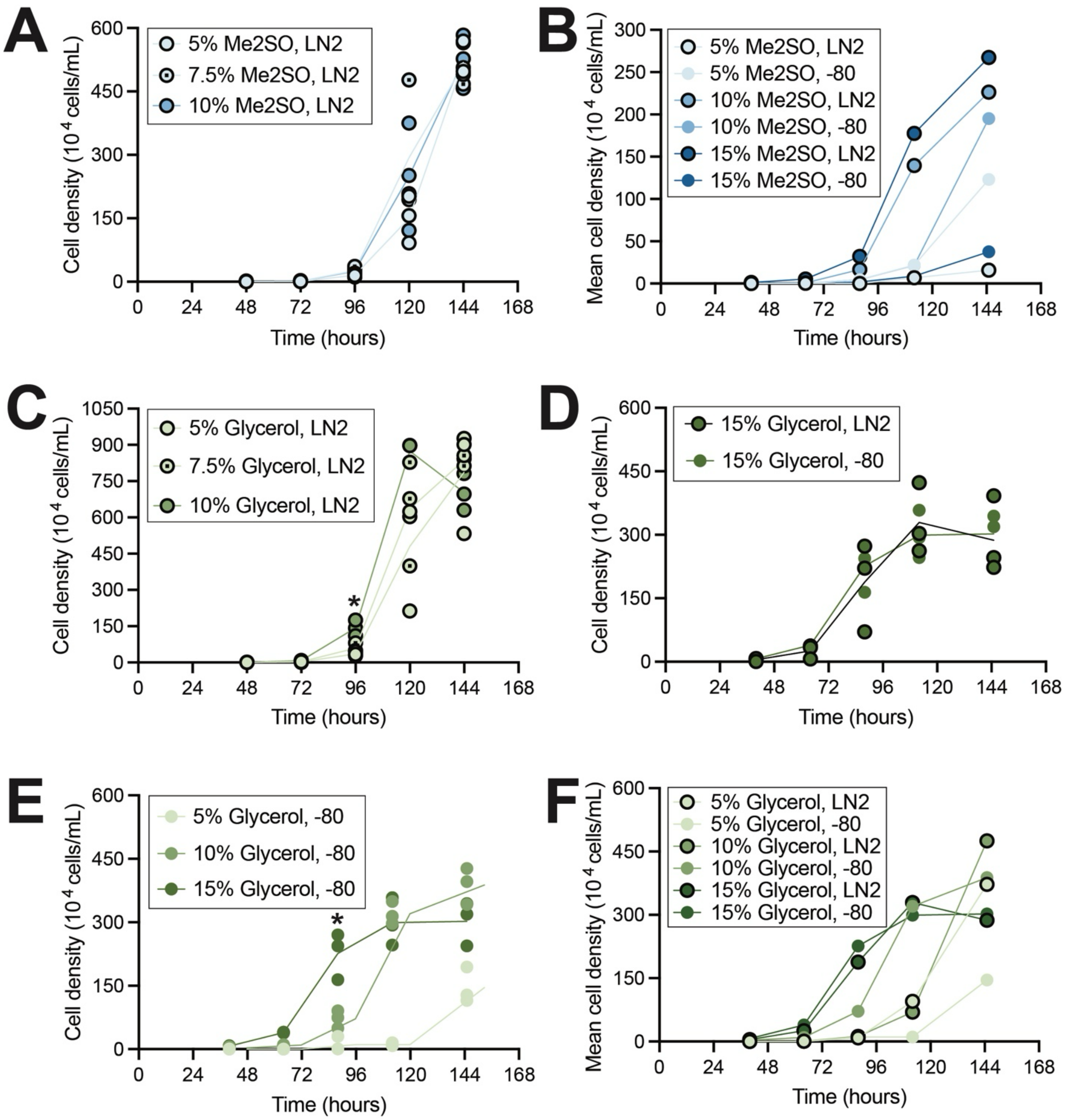


**Supplemental Figure 1. Additional testing of *S. rosetta* cultures confirmed 15% glycerol as the optimal cryoprotectant**. (A) *S. rosetta* cultures cryopreserved in 5%, 7.5%, or 10% DMSO and stored in LN2 did not show a significant difference in viability. (B) *S. rosetta* cultures cryopreserved in 5%, 10%, or 15% DMSO had higher mean cell densities when stored in LN2. At 145 hours, cell cultures cryopreserved in 15% DMSO and stored in LN2 had significantly higher viability than those stored at -80°C (*p=0.0294). For 10% DMSO, the higher mean cell density observed with LN2 storage was not significant. (C) At 96 hours, *S. rosetta* cultures cryopreserved with 10% glycerol showed better viability than with 5% glycerol (*p=0.0428). (D) *S. rosetta* cultures cryopreserved with 15% glycerol and recovered in high-nutrient media were equally viable between storage in LN2 or at -80°C. (E) *S. rosetta* cultures cryopreserved in 15% glycerol, stored at -80°C, and recovered in high-nutrient media had higher viability than those cryopreserved in 5% or 10% glycerol. At 88 hours, *S. rosetta* cultures cryopreserved in 15% glycerol showed the best viability after storage at -80°C (*p=0.0147 for 5% vs 15% glycerol; p=0.0283 for 10% vs 15% glycerol). (F) *S. rosetta* cultures cryopreserved with 15% glycerol had higher mean cell densities when stored in LN2 or at -80°C compared to cells cryopreserved with 5% or 10% DMSO, which had lower cell densities when stored in LN2 (in B). For all charts in Supplemental Figure 1, significance was determined by a two-way ANOVA multiple comparisons test with Geisser–Greenhouse and Šídák corrections (calculations and charts, Prism v. 10.1.1).

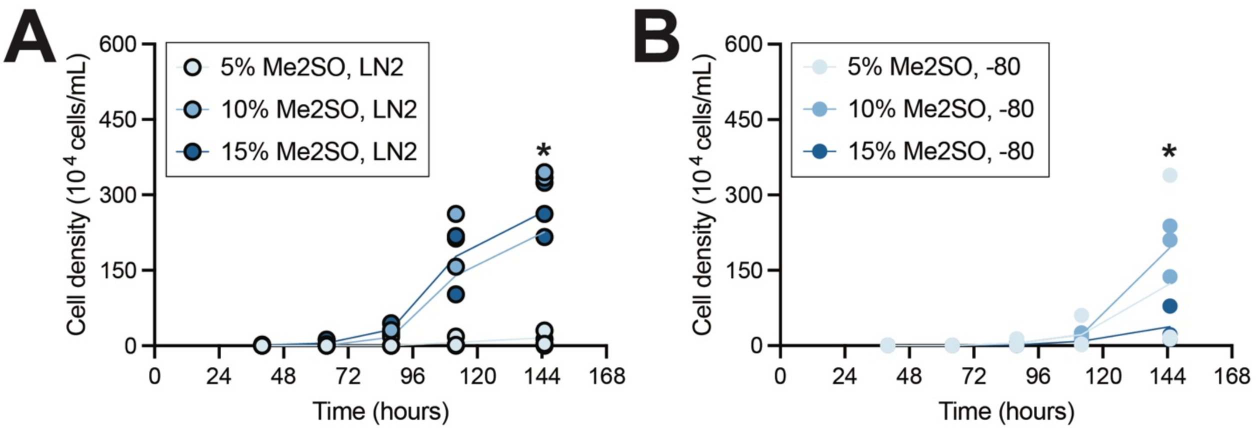


**Supplementary Figure 2. 10% DMSO and liquid nitrogen storage offer the ideal cryoprotectant percentage and storage conditions for cultures cryopreserved with DMSO**. (A) *S. rosetta* cultures preserved with DMSO, stored in LN2, and recovered in high-nutrient media were most viable when cryopreserved with at least 10% DMSO. At 145 hours, cell cultures cryopreserved with 5% DMSO were not as viable as those cryopreserved in 15% DMSO (*p=0.0215). Cultures cryopreserved in 10% DMSO were not significantly different than cultures cryopreserved in 15% DMSO (p=0.9368). (B) *S. rosetta* cultures preserved with DMSO, stored in an ultra-low temperature freezer (-80°C), and recovered in high-nutrient media were most viable when cryopreserved with 10% DMSO. At 145 hours, cell cultures cryopreserved with 10% DMSO were more viable than cells cryopreserved with 15% DMSO (*p=0.0341). Cultures cryopreserved in 5% DMSO were not significantly different in viability from cultures cryopreserved in 10% DMSO (p=0.8130) Significance was determined by a two-way ANOVA multiple comparisons test with Geisser–Greenhouse and Šídák corrections (calculations and charts, Prism v. 10.1.1).

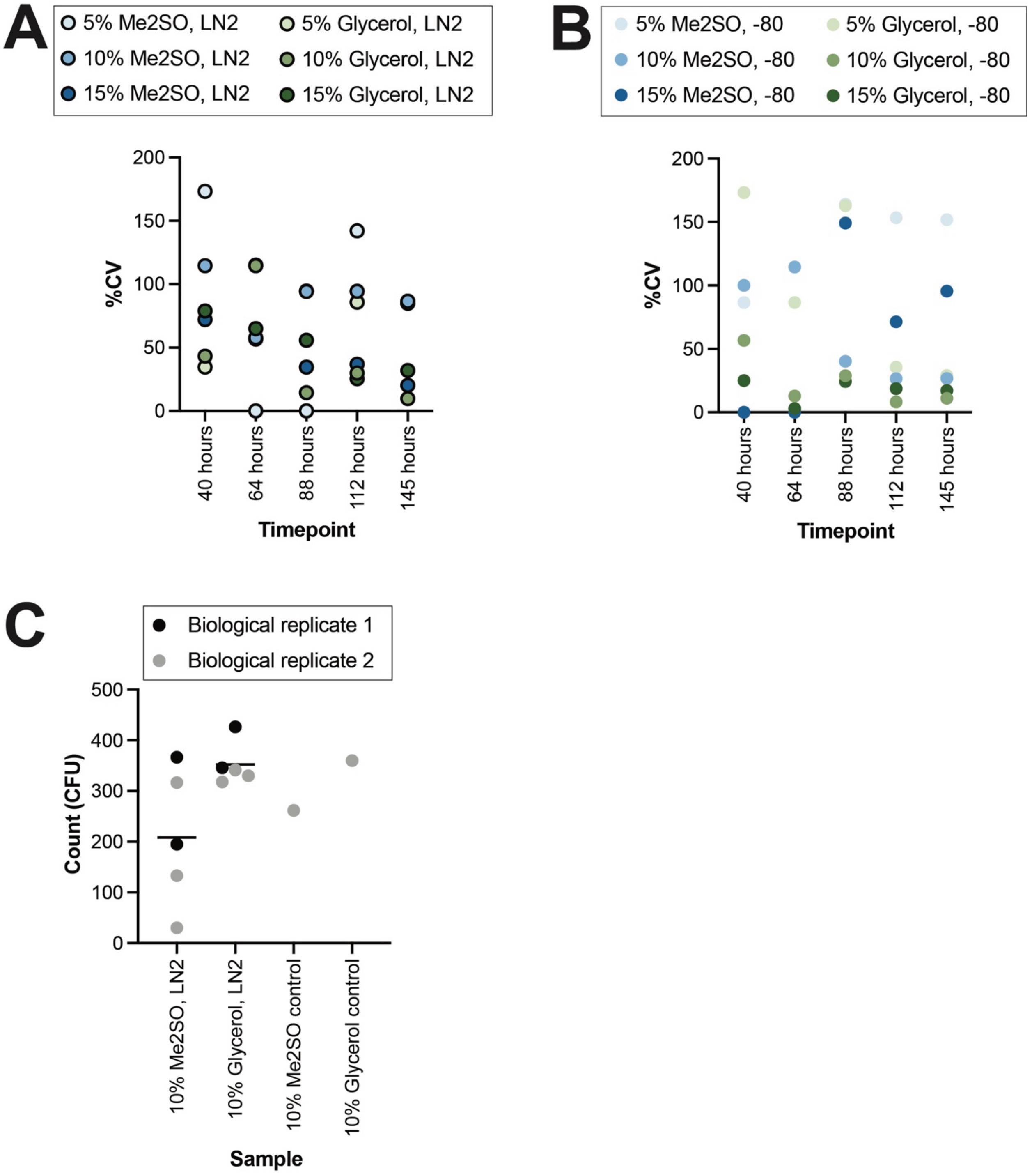


**Supplemental Figure 3. Additional testing of *S. rosetta* cultures confirmed 15% glycerol as the optimal cryoprotectant**. (A-B) *S. rosetta* cultures cryopreserved with 10% DMSO, the current cryoprotectant standard, showed more variability (%CV, coefficient of variation) at most timepoints when stored in LN2 (A) or at -80°C (B) compared to 10% or 15% glycerol. (C) Co-cultures of *S. rosetta* and *E. pacifica* yielded more *E. pacifica* colonies when cryopreserved in glycerol compared to DMSO. Bacterial colonies are given in CFU (colony-forming units), and individual replicates are represented by a single point (charts, Prism v. 10.1.1).
